# A Precision Medicine Approach to Stress Testing Using Metabolomics and Microribonucleic Acids

**DOI:** 10.1101/2020.02.06.936757

**Authors:** Alexander T. Limkakeng, Laura-Leigh Rowlette, Ace Hatch, Andrew B. Nixon, Olga Ilkayeva, David L. Corcoran, Jennifer L. Modliszewski, S. Michelle Griffin, Ephraim L. Tsalik, Geoffrey S. Ginsburg, Deepak Voora

**Affiliations:** Division of Emergency Medicine, Duke University, Durham, North Carolina, United States of America; Sequencing & Genomic Technologies Shared Resource, Duke Center for Genomic and Computational Biology, Duke University, Durham, North Carolina, United States of America; Division of Medical Oncology, Duke University, Durham, North Carolina, United States of America; Duke Molecular Physiology Institute, Duke University, Durham, North Carolina, United States of America; Genomic Analysis and Bioinformatics Shared Resource, Duke Center for Genomic and Computational Biology, Duke University, Durham, North Carolina, United States of America; Center for Applied Genomics & Precision Medicine, Duke University, Durham, North Carolina, United States of America; Emergency Medicine Service, Durham Veterans Affairs Health Care System, Durham, NC USA; Division of Infectious Diseases & International Health, Department of Medicine, Duke University, Durham, North Carolina, United States of America; Division of Cardiology, Duke University, Durham, North Carolina, United States of America

**Keywords:** Coronary heart disease, stress testing, metabolomics, transcriptomics, microRNA, precision medicine

## Abstract

**Background:** Acute coronary syndrome (ACS) is a growing global health problem, and precision medicine techniques hold promise for the development of diagnostic indicators of ACS. In this pilot, we sought to assess the utility of an integrated analysis of metabolomic and microRNA data in peripheral blood to distinguish patients with abnormal cardiac stress testing from matched controls.

**Methods:** We used prospectively collected samples from emergency department (ED) patients placed in an ED-based observation unit who underwent stress testing for ACS. We isolated microRNA and quantified metabolites from plasma collected before and after stress testing in patients with myocardial ischemia on stress testing versus those with normal stress tests. The combined metabolomic and microRNA data were analyzed jointly for case (ischemia) and 1:1 matched control patients in a supervised, dimension-reducing discriminant analysis. Two integrative models were implemented: a baseline model utilizing data collected prior to stress-testing (T0) and a stress-delta model, which included the difference between post-stress test (T1) and pre-stress test (T0).

**Results:** Seven case patients with myocardial ischemia on ED cardiac stress testing (6 females, 85% Caucasian, mean Thrombolysis In Myocardial Infarction Score=3, 4 patients ultimately received percutaneous coronary intervention) were 1:1 age and sex-matched to controls. Several metabolites and microRNAs were differentially expressed between cases and controls. Integrative analysis of the baseline levels of metabolites and microRNA expression showed modest performance for distinguishing cases from controls with an overall error rate of 0.143. The stress-delta model showed worse performance for distinguishing cases from controls, with an overall error rate of 0.500.

**Conclusions:** Given our small sample size, results are hypothesis-generating. However, this pilot study shows a potential method for a precision medicine approach to cardiac stress testing in patients undergoing workup for ACS.

## Introduction

Acute Coronary Syndrome (ACS). remains one of the most significant health problems globally(1). A cornerstone of risk assessment for ACS is provocative stress testing(2–5). Conceptually, stress testing is composed of two elements, a stressor and an evaluation of function. The evaluative function of stress testing currently depends on imaging technology to evaluate for the presence of ischemic myocardium. However, imaging evaluation requires specialized equipment and technical expertise. A blood-based biomarker approach to cardiac stress testing could obviate the need for expensive equipment or highly trained personnel.

MicroRNAs are small non-coding segments of RNA circulating that can be found in the bloodstream and act as paracrine regulators of local cellular gene transcription(6). MicroRNA profiles may be able to distinguish among various causes of myocardial injury, e.g. myocardial ischemia from heart failure(7). Based on prior literature, a number of microRNAs seem promising for identifying myocardial ischemia from coronary artery disease: miR-1(8–12), miR- 133(8, 9, 13), miR-208(9, 12, 14), miR-499(8, 9, 12–14), miR-126(10, 13, 15). Thus, microRNAs are a promising area of biomarker research.

Metabolomics is another promising modality for the characterization of function in high-energy utilization organs such as the heart. Such analyses examine a wide range of fundamental biological molecules, many of which are connected to underlying metabolic processes in the body such as fatty acids and oxidation products. The concentration of these molecules can also rapidly change in response to acute disease states. It has been shown that certain amino acids and acylcarnitines levels in peripheral blood are associated with long-term risk of cardiovascular disease, particularly coronary related(16–18). We previously reported the analysis of stress-induced changes in selected metabolites including amino acids and acylcarnitines(19).

In current clinical practice, the information gathered from stress testing is usually reduced down to a single data dimension to simplify decision-making. However, it is widely recognized that a precision medicine strategy for chest pain evaluation will require expanding the number of biomarkers (whether blood-based, imaging, or in other forms) and to integrate information to provide a more accurate answer. The challenge associated with expanding the number of biomarkers is that very large datasets can be problematic for biostatistical analysis. However, several dimension-reducing biostatistical approaches now allow the integration of large datasets from different categories of molecules(20).

In this paper, we used transcriptomic and metabolomic approaches to demonstrate the feasibility of a biomarker-based stress test using precision medicine techniques. We also sought to develop the technical capabilities and protocols to study serially measured microRNAs and metabolites in patients undergoing cardiac stress testing for symptoms of ACS. We believe this novel model of stress testing represents an exciting opportunity to apply a precision medicine approach to cardiac disease diagnosis and prognosis.

## Methods

### Study setting and population

We conducted a pilot study to determine whether serial microRNA and metabolomic data could be combined to enhance the diagnostic performance of cardiac stress testing. We used peripheral blood samples in EDTA collection tubes from a biorepository created to study changes in high-sensitivity troponin and B-type natriuretic peptide during stress testing. This biorepository has been previously described(21, 22). Briefly, samples were collected from adult emergency department (ED) patients who had symptoms of ACS and who underwent stress testing in our observation unit. All patients, as a condition of enrollment, underwent standard symptom-limited Bruce Protocol exercise echocardiogram tests as part of their usual care. These tests reported the presence or absence of inducible myocardial ischemia, defined as stress-induced regional wall motion abnormality in at least one segment. All tests were interpreted by board-certified cardiologists who were blinded to any biomarker data. Two reviewers independently confirmed the accuracy of the reports for this study. Patients had follow-up phone calls at one year.

### Metabolomic analyses

Similar to our prior work(19), we used standard mass spectrometry to determine plasma quantities of selected acylcarnitines and amino acids, as previously described (Table 1)(23). We used standard liquid-handling steps for the Genesis RSP 150/4 Robotic Sample Processor (Tecan AG, Maennedorf, Switzerland). Plasma samples were spiked with cocktails of stable isotope-labeled standards specific to each assay module for quantitative measurement of these targeted analytes. The proteins were precipitated with methanol, supernatant dried, and esterified with hot, acidic methanol (acylcarnitines) or n-butanol (amino acids). We then used tandem mass spectrometry on a Quattro Micro instrument (Waters Corporation, Milford, MA) to analyze acylcarnitines and amino acids. The lower level of quantitation for amino acids was 0.5 μM and for acylcarnitines the limit of quantitation was 0.015 μM.

**Table 1.**
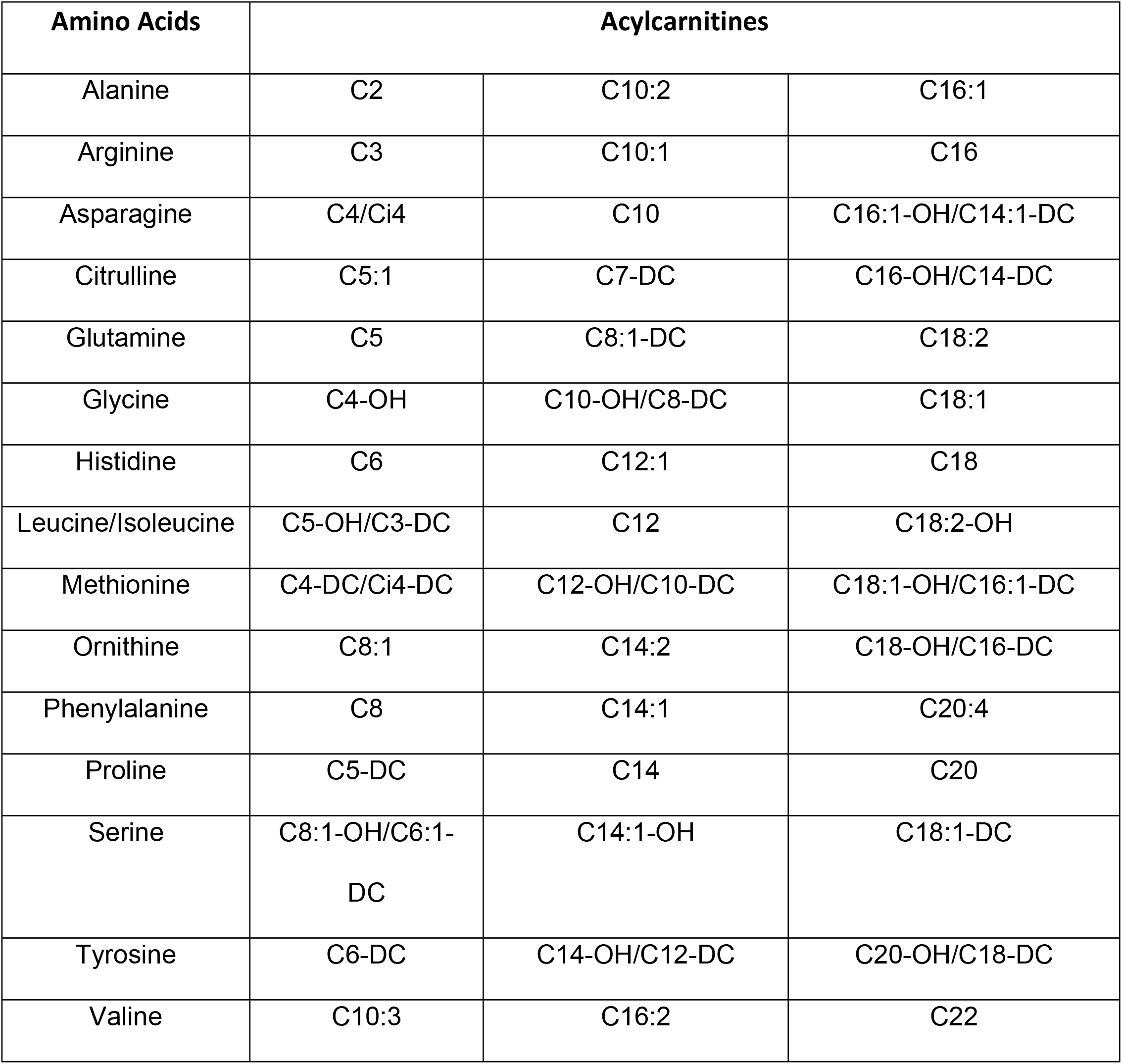
Metabolites Assayed

| Amino Acids | Acylcarnitines |  |  |
| --- | --- | --- | --- |
| Alanine | C2 | C10:2 | C16:1 |
| Arginine | C3 | C10:1 | C16 |
| Asparagine | C4/Ci4 | C10 | C16:1-OH/C14:1-DC |
| Citrulline | C5:1 | C7-DC | C16-OH/C14-DC |
| Glutamine | C5 | C8:1-DC | C18:2 |
| Glycine | C4-OH | C10-OH/C8-DC | C18:1 |
| Histidine | C6 | C12:1 | C18 |
| Leucine/Isoleucine | C5-OH/C3-DC | C12 | C18:2-OH |
| Methionine | C4-DC/Ci4-DC | C12-OH/C10-DC | C18:1-OH/C16:1-DC |
| Ornithine | C8:1 | C14:2 | C18-OH/C16-DC |
| Phenylalanine | C8 | C14:1 | C20:4 |
| Proline | C5-DC | C14 | C20 |
| Serine | C8:1-OH/C6:1-DC | C14:1-OH | C18:1-DC |
| Tyrosine | C6-DC | C14-OH/C12-DC | C20-OH/C18-DC |
| Valine | C10:3 | C16:2 | C22 |

### MicroRNA analyses

We extracted RNA using the Qiagen miRNeasy Serum/Plasma Advanced Kit (Qiagen, Frederick, MD) from plasma collected in ethylenediamine tetraacetic acid (EDTA) tubes. We used a standard QIASeq miRNA Library Kit protocol (Qiagen, Frederick, MD) for library preparation, and library quality control (QC) was performed on the Agilent Bioanalyzer with the Deoxyribonucleic Acid (DNA) High-Sensitivity Assay. Samples were sequenced on the Illumina (San Diego, CA) HiSeq 4000 Sequencer at 50 bp Single Read.

### Data Analysis

For all analytes, comparisons were made between cases and controls at baseline (baseline model), and the difference between pre- and post- stress test (stress-delta model).

#### MicroRNA analysis

SmRNA-seq data were processed using the Trim Galore toolkit(24), which employs Cutadapt(25) to trim low-quality bases and Illumina sequencing adapters from the 3’ end of the reads. Only reads that were 18-28 nucleotides in length after trimming were kept for further analysis. Reads were mapped to the hg19 version of the human genome using the Bowtie alignment tool(26). Reads were kept for subsequent analysis if they mapped to no more than 13 genomic locations. Gene counts were compiled using custom scripts that compare mapped read coordinates to the miRbase microRNA database(27). Reads that match the coordinates of the known mature microRNAs were kept if they perfectly matched the coordinates of the miRNA seed while not varying by more than 2 nucleotides on the 3’ end of the mature miRNA. Only mature miRNAs that had at least 10 reads in any given sample were used in subsequent analysis. Normalization was performed using the DESeq2 Bioconductor package from the R statistical programming environment applying the ‘poscounts’ approach to eliminate systematic differences across the samples(28). The normalized data were log-transformed and differential expression was tested using linear regression. For the stress-delta model, we employed a mixed-effects model with the patient ID as a random effect. The false discovery rate was used to adjust for multiple hypothesis testing.

Targeted metabolite data were log-transformed prior to analysis and a PCA was conducted to assess for the presence of outliers and confounding demographic factors. MicroRNA-Seq count data were also log-transformed. For the integrative analysis, microRNAs that were missing in half or more of the samples were removed from the data set. All integrative analyses were conducted with baseline (“T0”, pre-stress test) and delta (“T1” – “T0”) data sets, where T1 corresponds to post–stress test samples.

#### Regularized Canonical Correlation Analysis

Regularized canonical correlation analysis (rCCA) seeks to extract latent variables that maximize the correlation between the two data sets, but with an additional regularization step that reduces the number of variables contributing to each component. An initial leave-one-out cross-validation step can be performed to select the regularization parameters for each data set. To explore correlation between the metabolomic and microRNA baseline and delta datasets, a rCCA was performed in the mixOmics package in R. Five components were retained in our final model.

#### Integrative Sparse Discriminative Analysis

To identify metabolites and microRNAs that discriminate between control and case subjects, we examined both datasets jointly in the mixOmics package in R using the DIABLO (Data Integration Analysis for Biomarker discovery using Latent variable approaches for ‘Omics studies) method. DIABLO is a supervised, dimension-reducing discriminant analysis using a sparse projection to latent structures analysis with a discriminant component; this performs similar to a canonical correlation analysis with the exception that covariance rather than correlation is maximized.

After determining the optimal number of components, the number of variables for each component was chosen through leave-one-out cross-validation over a grid of possible number of variables per component (minimum=1, maximum=50 and 30 for the baseline and delta data sets, respectively). Performance of the final model was assessed with leave-one-out cross-validation with the centroids distance.

## RESULTS

The baseline demographic and clinical characteristics of each subject are summarized in Table 2. Patients had a high rate of hypertension, hyperlipidemia, and diabetes. Six of the 7 Case patients had subsequent coronary angiography during the index visit, with 5 of them having at least one artery with stenosis >50%. Four patients underwent subsequent percutaneous coronary interventions. All Control patients underwent follow up at 1 year from their index ED visit without any cardiac diagnosis being made.

**Table 2.** Patient Demographics and Clinical Characteristics

| Characteristic | Cases<br>N (%) | Controls<br>N (%) |
| --- | --- | --- |
| Age (Years), Mean (Range) | 63.9 (54-76) | 64.0 (55-71) |
| Sex |  |  |
| Male | 1 (14.3%) | 1 (14.3%) |
| Female | 6 (85.7%) | 6 (85.7%) |
| Race |  |  |
| Black or African American | 1 (14.3%) | 3 (42.9%) |
| White / Caucasian | 6 (85.7%) | 4 (57.1%) |
| Hypertension | 6 (85.7%) | 4 (57.1%) |
| Diabetes | 4 (57.1%) | 3 (42.9%) |
| History of Tobacco Use | 3 (42.9%) | 0 (0%) |
| Hyperlipidemia | 6 (85.7%) | 3 (42.9%) |
| Past Myocardial Infarction | 4 (57.1%) | 1 (14.3%) |
| History of Coronary Artery Disease | 5 (71.4%) | 2 (28.6%) |

### Metabolomics Results

The concentrations of assayed amino acids and acylcarnitines both before and after stress testing for each subject are available in S1 File. Among acylcarnitines, acetylcarnitine (C2) and the hexanoylcarnitine (C6-DC/C8-OH) showed the greatest absolute differences in baseline values between cases and controls (7.54 μM versus 9.92 μM (p=0.54) and 0.12 μM versus 0.08 μM, (p=0.16) respectively). Among amino acids, glycine, proline, and tyrosine showed the greatest absolute differences in baseline values between cases and controls (421.04 μM versus 332.96 μM, (p=0.17); 184.87 μM versus 222.92 μM, (p=0.42); and 62.24 μM versus 75.08 μM, (p=0.13), respectively).

The greatest absolute differences in stress-delta values between cases and controls were seen in metabolites octanoylcarnitine (C8) and decanoylcarnitine (C10) (−0.004 μM versus −0.06 μM (p=0.39) and −0.02 μM versus −0.10 μM, (p=0.42) respectively). Among amino acids proline, valine, and asparagine showed the highest stress delta greatest absolute differences in stress-delta values between cases and controls (33.86 μM versus −0.79 μM, (p=0.24); 4.80 μM versus −21.75 μM, (p=0.44); and 10.80 μM versus −15.57 μM, (p=0.26).

### MicroRNA Results

Among 1,238 microRNAs, 52 were differentially expressed (p<0.05) between cases and controls at baseline and 12 had significantly different stress-deltas with unadjusted analysis. These microRNAs are listed in S2 File. We constructed heat maps (Figs 1 and 2) to show differential baseline and stress-delta microRNA expression in cases and controls. PCA plots demonstrated that moderate variance was explained by MicroRNA principal components (S1-S3 Figs).

**Fig 1.**
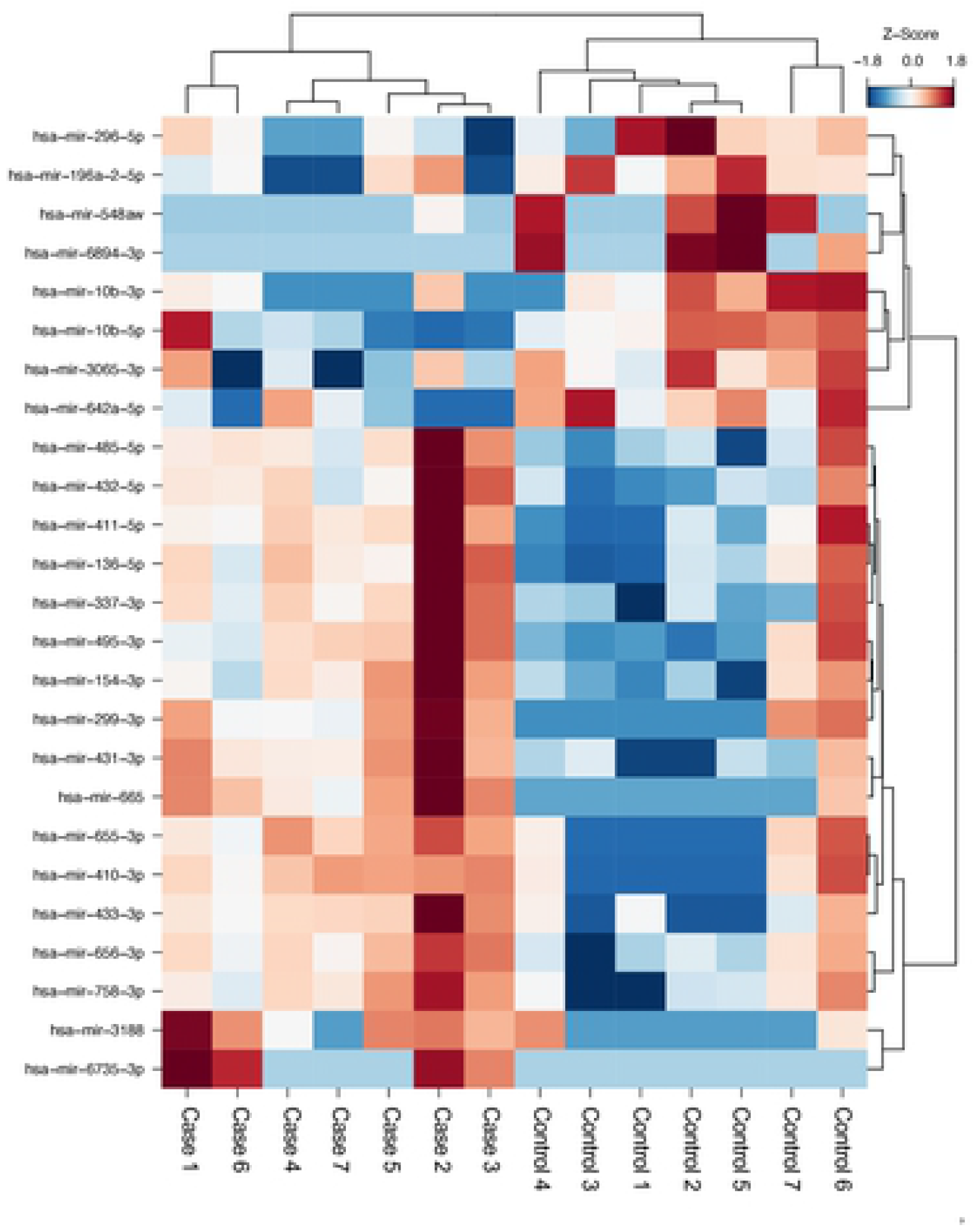
Baseline and Stress-Delta MicroRNA Heatmap. Heat maps demonstrating key baseline and stress-delta microRNAs that are different between case (myocardial ischemia) and matched control patients. The Baseline model (Fig 1) shows the z-score transformed expression value and the Stress-Delta heatmap (Fig 2) shows the log2 (fold-change) values for each patient across time. Both heatmaps have been clustered by both genes and samples using a correlation distance with complete linkage.

**Fig 2.**
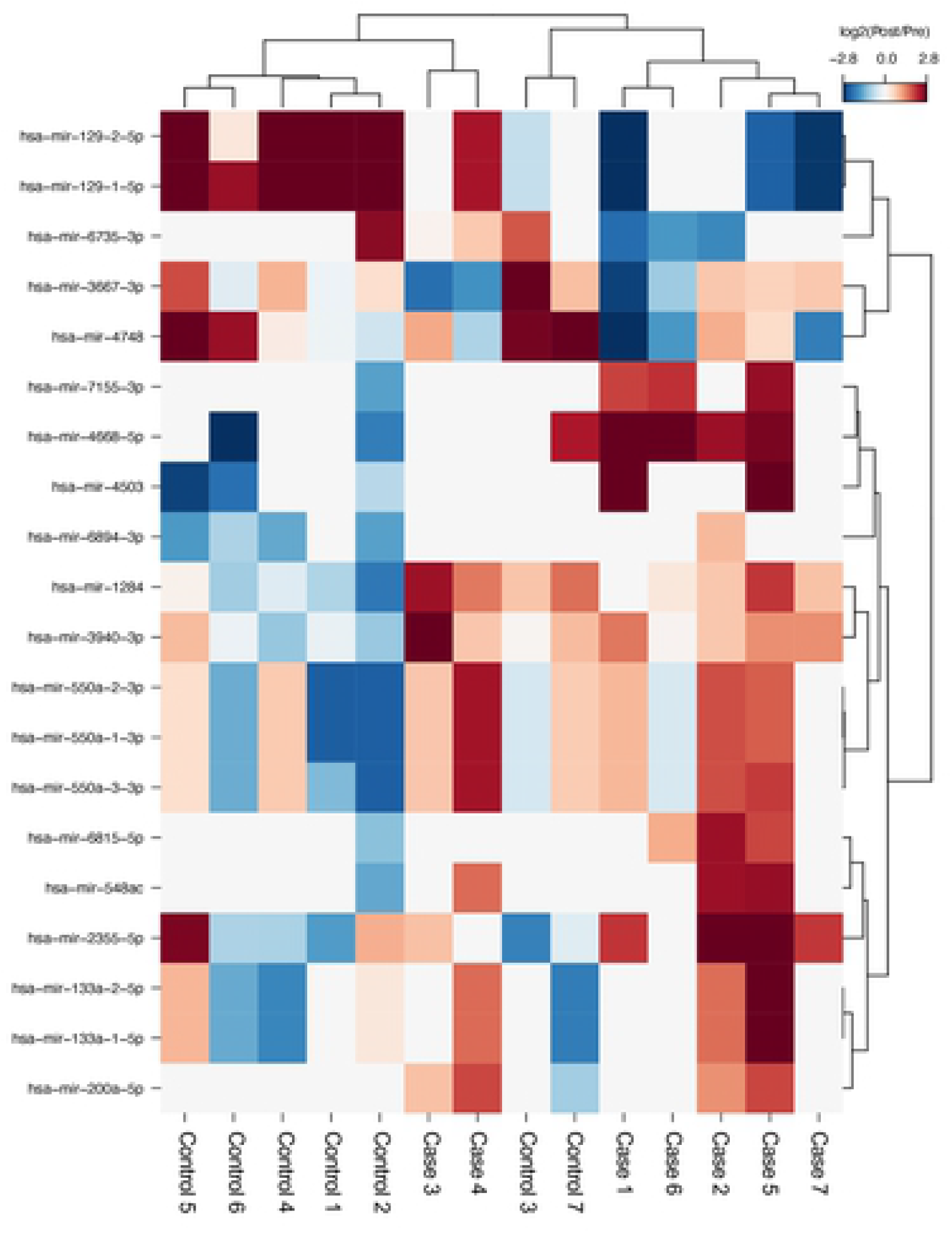
Baseline and Stress-Delta MicroRNA Heatmap. Heat maps demonstrating key baseline and stress-delta microRNAs that are different between case (myocardial ischemia) and matched control patients. The Baseline model (Fig 1) shows the z-score transformed expression value and the Stress-Delta heatmap (Fig 2) shows the log2 (fold-change) values for each patient across time. Both heatmaps have been clustered by both genes and samples using a correlation distance with complete linkage.

### Integrative Analysis

We performed rCCA to assess the correlation structure of the metabolomics and microRNA data. Figs 3 and 4 shows the baseline and stress-delta correlations of microRNAs and metabolites derived from the rCCA analysis. Only 6 of the baseline and 18 of the 64 stress-delta metabolites showed correlations above 0.65 with specific microRNAs, suggesting that combining the two datasets provided additive information.

**Fig 3.**
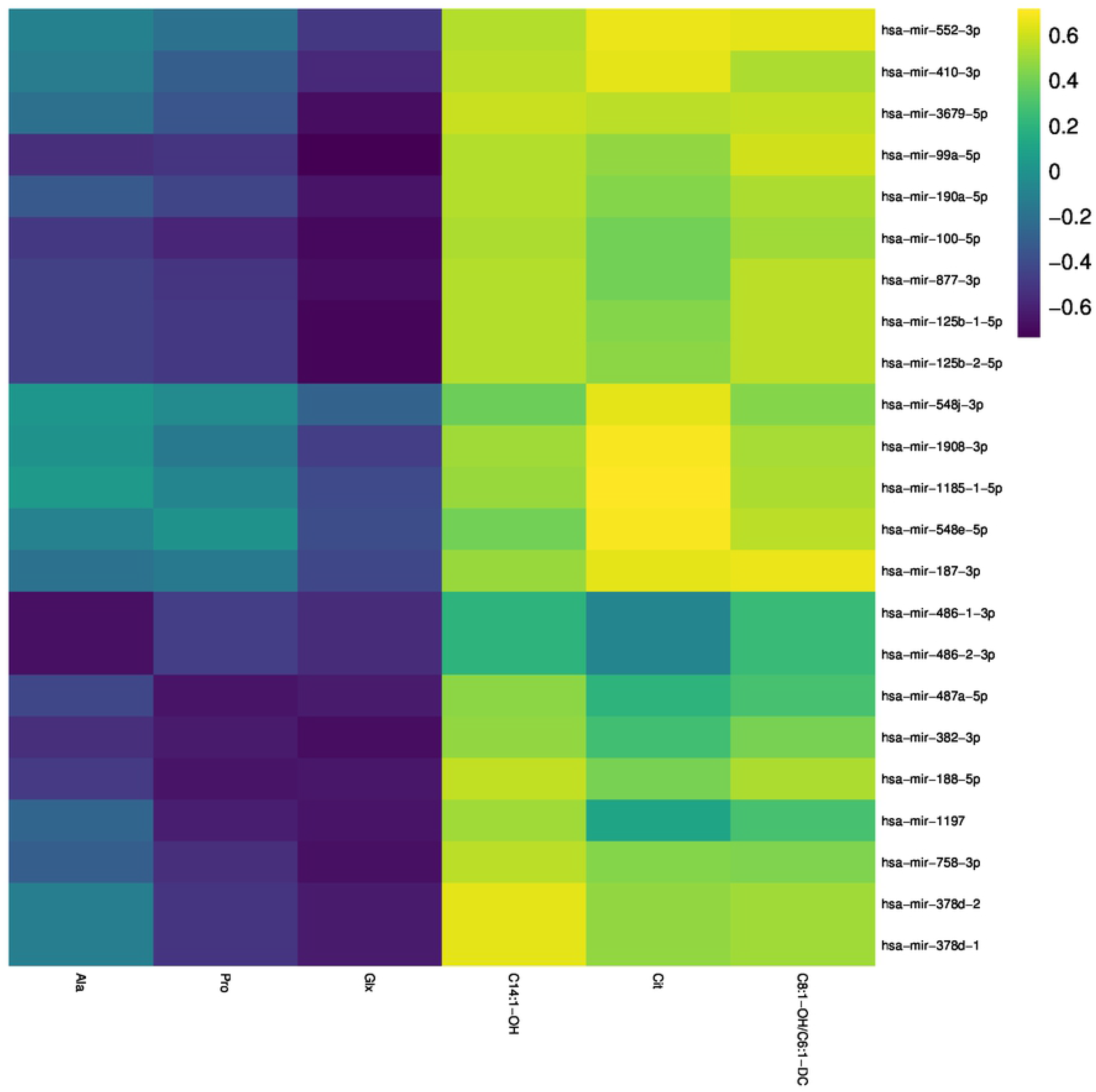
Regularized Canonical Correlation Analysis Heat Map of MicroRNAs and Metabolites. Regularized Canonical Correlation Analysis heatmap showingcorrelations between baseline (3) and stress-delta (4) microRNAs and metabolites as a result of stress testing.

**Fig 4.**
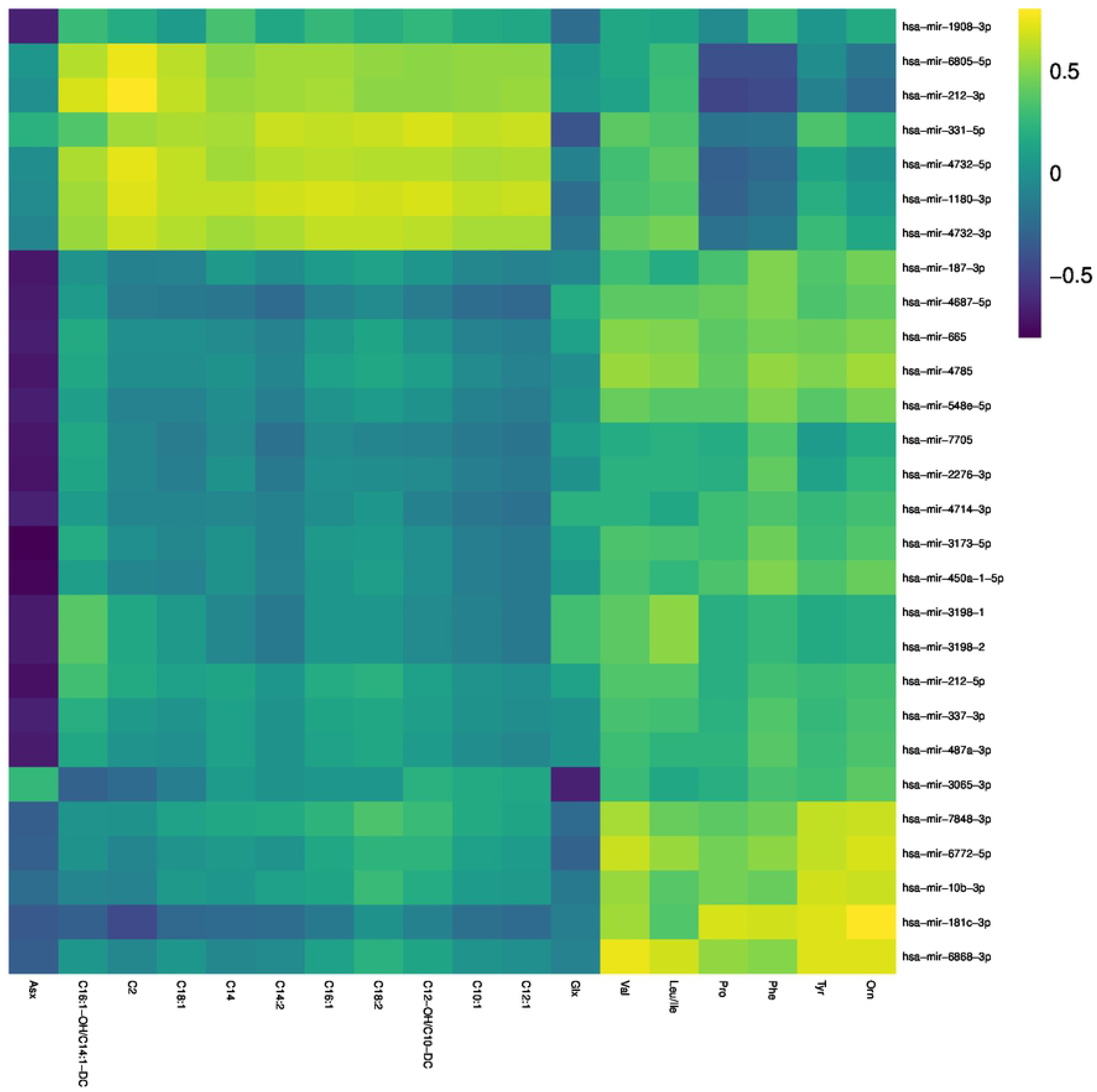
Regularized Canonical Correlation Analysis Heat Map of MicroRNAs and Metabolites. Regularized Canonical Correlation Analysis heatmap showing correlations between baseline (3) and stress-delta (4) microRNAs and metabolites as a result of stress testing.

For the integrative discriminant analysis, both the baseline model and the stress-delta model produced a single latent component. We calculated error rates for our integrated analysis model for predicting cases or controls. Integrative analysis of metabolite levels and microRNA expression at baseline showed modest performance for distinguishing cases from controls, with an overall error rate of 0.143 (Table 3). Using stress-delta data actually led to a worse error (0.500) for distinguishing cases from controls.

**Table 3.** Integrative Model Performance. Error rates are shown for each group individually (Case and Control). The weighted vote error rate is calculated by assigning each data set a weight based on the correlation between its latent component and the outcome.

| Group | Weighted Vote Baseline | Weighted Vote Stress Delta |
| --- | --- | --- |
| Case | 0.000 | 0.571 |
| Control | 0.286 | 0.429 |
| Overall Error Rate | 0.143 | 0.500 |

Figs 5 and 6 show a plot of individual subject scores for microRNA and metabolites latent components, using only those microRNAs and metabolites that were retained in the baseline and stress-delta integrative model. Cases and controls can be visually separated along these two axes representing each category’s latent component. It should be noted that these results represent a best-case estimate of our model’s ability to distinguish cases from controls given that the model was tested on the same data that it was trained on.

**Fig 5.**
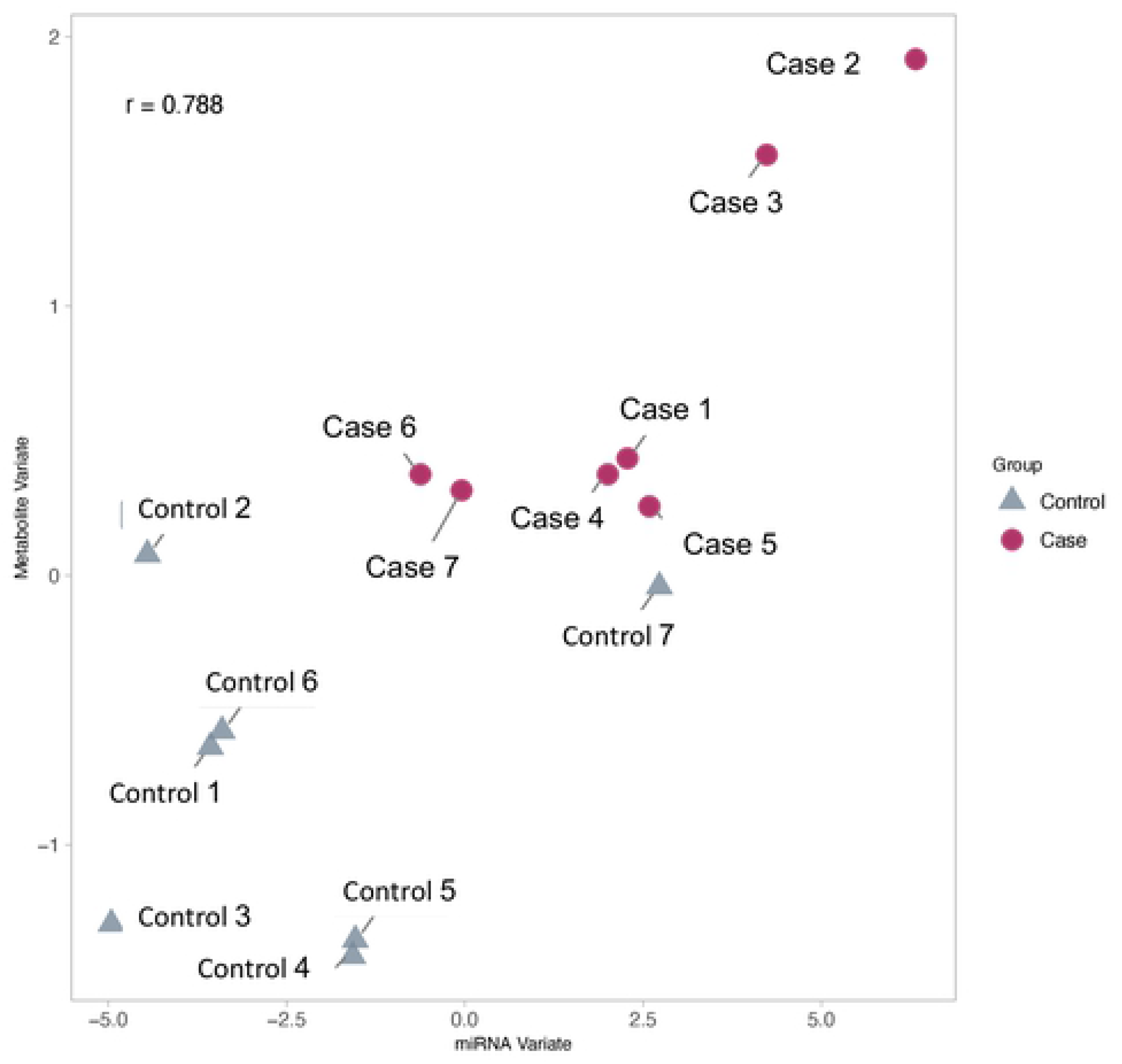
Plot of Coordinates for Individual Patients from the Baseline (Fig 5) and Stress-Delta (Fig 6) Integrative Model Using Both Metabolomic and MicroRNA Data. Control (grey triangle) vs. case (red circle) separation in the stress-delta integrative model.

**Fig 6.**
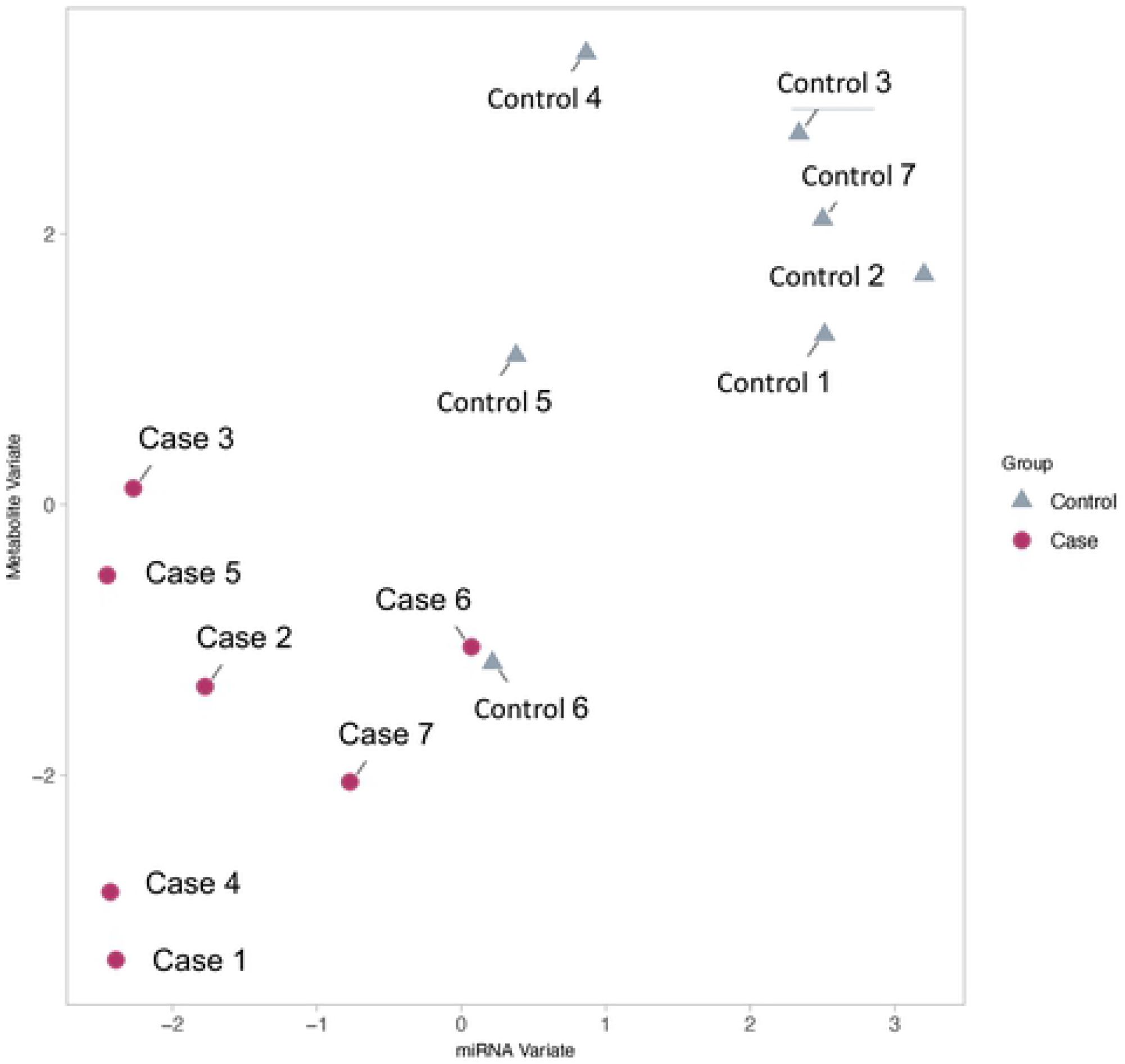
Plot of Coordinates for Individual Patients from the Baseline (Fig 5) and Stress-Delta (Fig 6) Integrative Model Using Both Metabolomic and MicroRNA Data. Control (grey triangle) vs. case (red circle) separation in the stress-delta integrative model.

Finally, we present the results for loadings to the model. These metabolites and microRNAs are the ones that were the most influential on the latent component. For the baseline model, the two analytes with the highest loadings were mir-665 and C18:1-DC. The loadings of selected metabolites and microRNAs for the component in the stress-delta model are shown in Fig 7. Analytes with the highest loadings are relatively equally divided between metabolites and microRNAs, suggesting value in combining both datasets.

**Fig 7.**
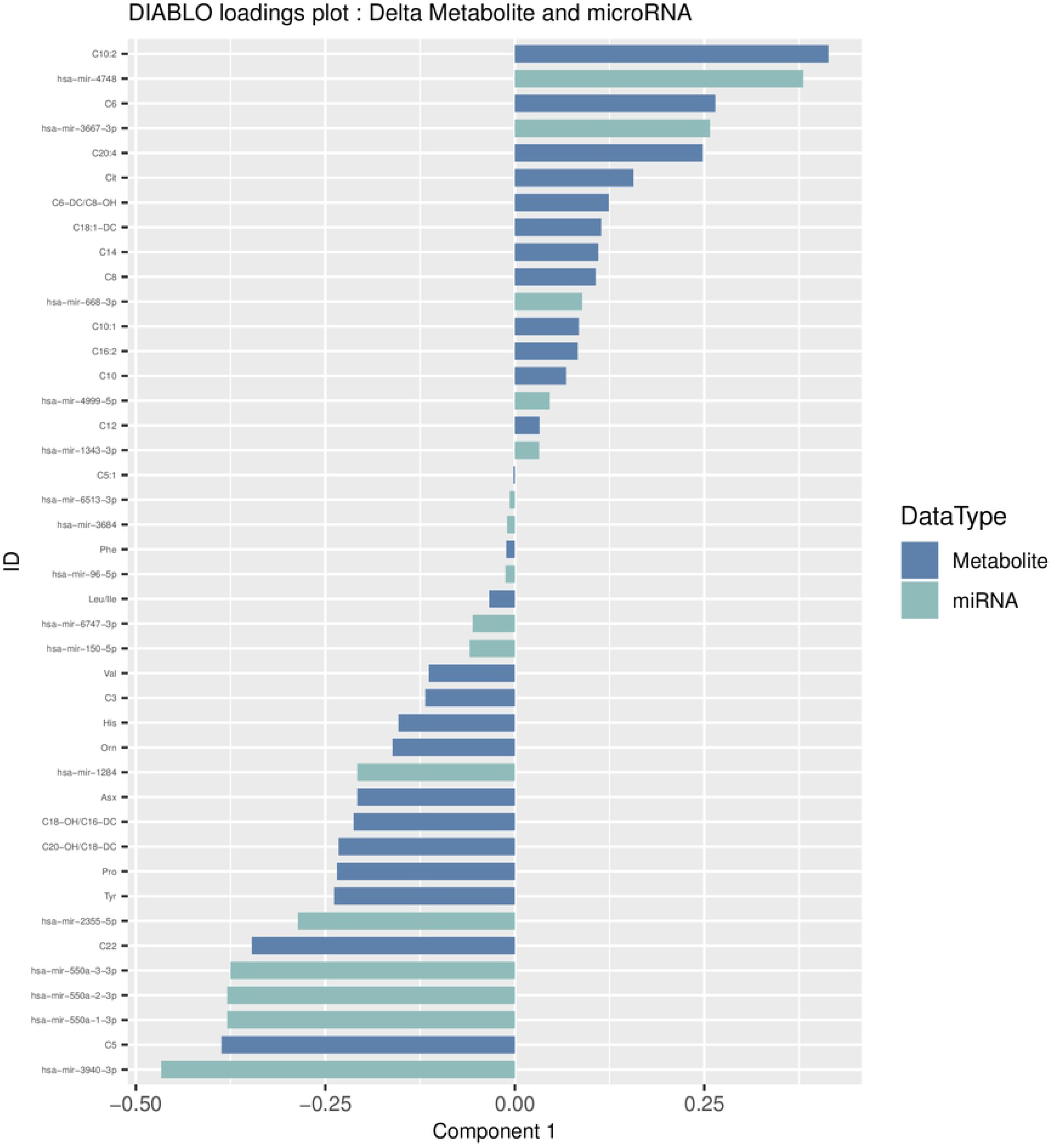
Metabolite and MicroRNA Loadings from the Latent Component Of Stress-Delta Integrative Model. Specific stress-delta metabolites and microRNAs that loaded to the latent component of the stress-delta model is shown. Case samples cluster on the lower end of the first component. A negative loading for a particular analyte indicates it increased in cases, whereas a positive loading indicates that is increased in controls.

## DISCUSSION

Although the observable features of myocardial ischemia have been noted for almost a century(29), stress testing has largely remain unchanged for the past three decades(5). Current stress test modalities do not widely utilize multiple modes of information to enhance prediction accuracy. Simple examples of multi-modal stress tests exist, e.g. the Duke Treadmill Score(30), which combines exercise tolerance information and electrocardiogram characteristics to make a prediction of future risk. However, these multi-modal stress tests do not take advantage of the large amounts of data that we are currently capable of collecting from patients’ blood samples. Although our current modalities of stress testing are sensitive for obstructive coronary disease, their accuracy can be further improved, particularly for identifying specific high-risk phenotypes that benefit from emerging therapies.

In contrast to the blood-based biomarker approach for evaluation of acute myocardial infarction, currently assessment for myocardial ischemia and/or obstructive coronary artery is heavily imaging-dependent. Use of serial biomarkers is not routine practice for assessing myocardial ischemia, especially in the context of a stress test. Thus, this study presents a novel paradigm for assessing patients for myocardial ischemia. Our current ability to serially measure multiple blood-based molecules presents an opportunity to develop more sophisticated multi-modal stress tests that incorporate large amounts of data.

We have previously examined the utility of blood-based biomarkers to enhance cardiac stress testing(19, 21, 22). In this pilot study, we outline the methodology to further develop a biomarker-augmented stress test using a precision medicine approach. First, we identified several differences at baseline, post-stress, and stress-delta between cases and controls, largely as a consequence of the large number of analytes we assessed. While a great deal of prior literature has examined baseline (resting) biomarkers for prediction of coronary heart disease, stress-delta biomarker assessments give us the ability to assess acute changes in response to a controlled ischemic event, with the benefit of within-patient control for baseline values. We were able to assess a large number of potential biomarkers in each blood sample, creating the possibility of a systems biology approach to biomarker discovery.

Systems biology is an efficient approach to both understanding pathophysiologic mechanisms and identifying clinically useful biomarkers. MicroRNAs are an ideal clinical biomarker target to identify myocardial ischemia because their peripheral blood concentration can change rapidly in response to disease and they remain stable and detectable in the peripheral bloodstream. Our study demonstrates that microRNAs can be easily isolated from peripheral blood plasma and analyzed accurately in serial fashion. Furthermore, many candidate microRNAs appear to differentiate patients with myocardial ischemia from those with normal studies, suggesting promise for future studies in larger patient cohorts.

Likewise, metabolomics may be an ideal means to obtain information on the viability of the heart because of the organ’s dynamic nature. Numerous studies have demonstrated that resting baseline metabolite abnormalities are associated with adverse cardiovascular outcomes(18, 31, 32). Furthermore, myocardial ischemia is known to cause dysregulated energy utilization of myocardial cells(33). A previous study(17) demonstrated that a number of metabolites change dynamically in patients with ischemic stress tests compared to normal controls. Our prior work showed that alanine, C14:1-OH, C16:1, C18:2, C20:4 demonstrated patterns of acute changes in ischemic patients that were different from normal controls. In the current study, our small sample size was underpowered to confirm or refute this finding. However, combining metabolomics with microRNA data did provide additive diagnostic information. Other categories of molecules could be used in a precision medicine stress test, such as proteomics, immune mediators, catecholamine levels, and traditional markers of cardiac necrosis or stress.

As a pilot study, there are many limitations to this analysis. The small sample size and large number of analytes examined precludes us from making definitive statements about the importance of any specific analyte for our stress-delta paradigm. It is important to note that the error rates represent a best-case estimate of what is expected given that they are based on the same data the model was trained on. Our limited sample size prevented us from performing a validation in an independent cohort. In the future, we hope to collect a large set of samples which will enable us to have separate testing and validation cohorts to fully measure the robustness of this model. Furthermore, our patients were chosen from a cohort of patients referred for stress testing in a single center’s emergency department observation unit. Use of a biomarker-augmented stress test needs to be studied in a more representative patient sample in the future.

## CONCLUSIONS

In this pilot study of patients undergoing cardiac stress testing, we analyzed serially drawn blood samples for microRNA and metabolite levels. We demonstrated how these data could be used to differentiate patients with myocardial ischemia on imaging from normal controls. Based on this pilot, we intend to further study this paradigm of stress testing in a larger cohort. Our current paradigm of cardiac stress testing can be enhanced by systematic molecular profiling techniques. Future work should be conducted to identify the specific modalities and/or analytes that change dynamically in the setting of induced myocardial ischemia.

## Acknowledgements

We would like to acknowledge manuscript preparation assistance by Ms. Ashley Morgan.

The dataset(s) supporting the conclusions of this article is(are) included within the article (and its additional file(s)).

- Competing interests: Dr. Ginsburg report having an unlicensed patent on a metabolomic finding. Dr. Ginsburg also serves on the Scientific Advisory Board for CardioDx. Drs. Ginsburg, Limkakeng, and Voora have received research funding from Abbott Laboratories. Dr. Limkakeng has also received prior research grants from Roche International, and Siemens Healthcare Diagnostics. No other competing interests to declare.
- Funding: The current study was made possible by funding support via a Pilot Grant from the Emergency Medicine Foundation. Samples were collected in a prior study that was supported by an investigator-initiated grant from Abbott Laboratories. The authors retained possession of all data and decision on whether to publish at all times.

## Supporting information

**S1 File. Concentrations Of Assayed Amino Acids And Acylcarnitines Both Before And After Stress Testing.**

**S2 File. Differentially expressed MicroRNAs between Cases and Controls at Baseline and Stress Delta.**

**S1 Fig. Pre-Stress Test Principal Component Analysis Plot for MicroRNAs.** Principal component analysis of microRNA concentrations before stress testing to differentiate between case (myocardial ischemia) and matched control patients.

**S2 Fig. Post-Stress Test Principal Component Analysis Plot for MicroRNAs**. Principal component analysis of microRNA concentrations after stress testing to differentiate between case (myocardial ischemia) and matched control patients.

**S3 Fig. Principal Component Analysis Plot for MicroRNAs Pre-Stress Test Versus Post-Stress Test**. Principal component analysis of microRNA concentrations from before stress testing and after stress testing.

**S4 Fig. Receiver Operator Curve for Metabolites.**Stress-delta metabolite concentrations showed strong discriminative abilities individually. These results represent best case scenarios due to lack of validation cohort.

**S5 Fig. Receiver Operator Curve for MicroRNA.**Stress-delta MicroRNA concentrations showed strong discriminative abilities individually. These results represent best case scenarios due to lack of validation cohort.

